# Laser Speckle Contrast Imaging of Hepatic Microcirculation

**DOI:** 10.1101/2024.12.24.629390

**Authors:** Oleg Zhukov, Dmitry D. Postnov, Kamilla H. Hejn, Kim Ravnskjær, Olga Sosnovtseva

## Abstract

The liver controls blood homeostasis and depends critically on adequate blood supply. While the global regulation of liver blood flow via the hepatic arterial buffer response is well established, the mechanisms governing hepatic sinusoidal hemodynamics remain elusive. We use laser speckle contrast imaging to investigate the hepatic microvascular blood flow in anesthetized rats. Laser speckle contrast imaging offers a spatial resolution of a few micrometers, enabling visualization of individual microvessels, and a temporal resolution sufficient to track flow dynamics. This allowed us to resolve individual sinusoids and venules on the liver surface and to detect a reduction of the blood flow following local Angiotensin-II injections. We show that the blood flow oscillates with frequencies within the range of 0.05–0.4 Hz, which may be linked to rhythmic contraction of upstream blood vessels. Our findings provide insights into vessel-specific liver microcirculation *in vivo*, offering new opportunities to explore vascular dysfunction mechanisms in metabolic liver diseases.

## 1 Introduction

The liver is a major metabolic organ, which has a unique vascular system [1]. The liver receives about 25 % of the cardiac output via a dual blood supply consisting of oxygenated blood from the hepatic artery and nutrient-rich blood from the portal vein [2, 3]. Inside the liver, the terminal hepatic arterioles and terminal portal venules merge forming the sinusoids, i.e. the capillary bed of the liver [4–6]. The sinusoids feed the nuetrients and oxygen to the hepatocytes that detoxify the blood, control its glucose and fat levels, and synthetize proteins [1, 7]. The sinusoids then form the terminal central venules that drain the blood from the liver via the hepatic vein [8].

The liver is sensitive to hemodynamics in both arterial and venous blood flow. The hepatic arterial buffer response balances the two [2, 9]: when the portal venous flow decreases, the hepatic arteriolar flow increases via vasodilation, maintaining a proper oxygenation and a steady blood flow in the liver [10, 11]. Endogenous vasoactive molecules, such as Angiotensin II, influence the blood flow in the portal venules [12, 13], which dominate the total liver blood inflow [2] and differs from other veins by its stronger and rhythmic contractility [14–16]. The sinusoidal blood flow has been characterized as highly heterogenous [17, 18] and sensitive to sinusoidal contractions [19, 20]. However, little is known about intravital hemodynamics of the sinusoids.

Laser speckle contrast imaging (LSCI) is a rapidly developing wide-field imaging technique that quantifies blood flow velocity by measuring local changes in the light scattering produced by moving red blood cells [21–25]. LSCI has been instrumental in understanding microcirculation in brain [26, 27], kidney [28–31], and other organs [23, 32]. However, LSCI application in the liver remains limited to global flow measurements [33–35].

We demonstrate that LSCI can resolve individual liver microvessels, including the sinusoids, reliably measure the blood flow responses to vasoactive agents, and detect spontaneous oscillations of the microvascular blood flow.

## 2 Results

### 2.1 LSCI resolves individual hepatic microvessels

To quantify the blood flow changes in individual microvessels of the liver we performed laser speckle contrast imaging (LSCI) (Fig. 1a-d). Spatial maps of the temporal blood flow index (tBFI), generated after liver stabilization and averaging the tBFI over time, enabled visualization of blood vessels as small as 8 *μ*m in diameter (Fig. 1e, Fig. S1,S2) — a level of detail not achieved in previous LSCI studies of the liver [33–35].

**Figure 1:**
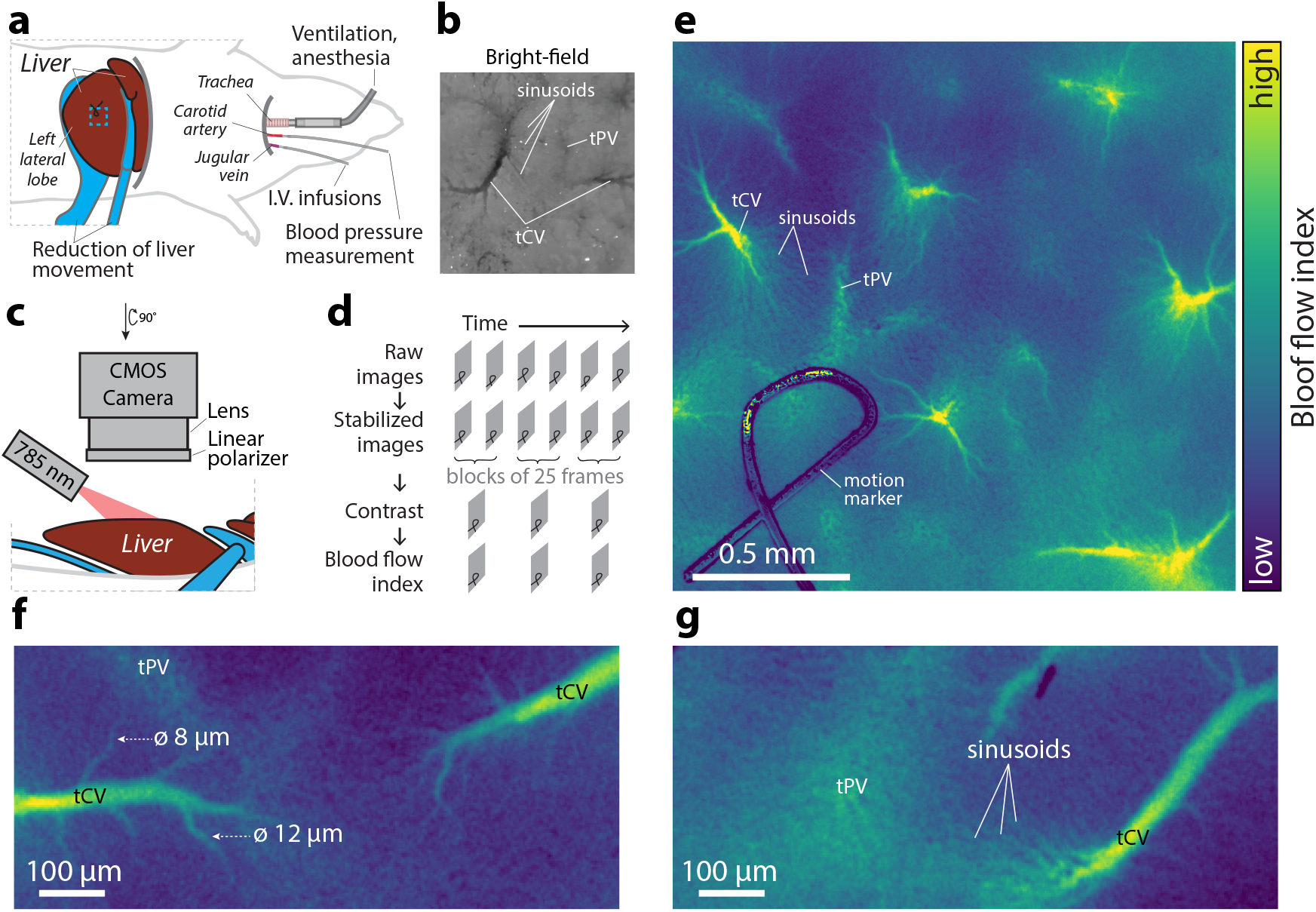
Experimental setup for imaging the hepatic microvasculature *in vivo*. **a**) Rat preparation viewed from above. Images were acquired from a region (blue rectangle) of the left lateral lobe of the liver. **b**) A bright-field image of the liver surface vasculature: terminal central venule (tCV), terminal portal venule (tPV), and sinusoids. **c**) Laser speckle imaging setup. The liver surface was illuminated with a 785 nm laser diode; the scattered light was captured by a CMOS camera (see Methods). **d**) Raw images were first stabilized by matching the position of the motion marker. The temporal laser speckle contrast and the blood flow index (tBFI) were calculated from the stabilized images. Blood flow index was averaged in time to obtain blood flow index maps. **e**) A tBFI map showing hepatic microvessels. See Fig. S1 for a tBFI map obtained from a larger region of the liver. **f,g**) Close-up of the terminal central venules (f) and sinusoids (g). See Fig. S2 showing sinusoids in another rat.

The blood vessels depicted in the tBFI maps closely resemble those observed through intravital (Fig. 1b) [10, 36, 37] and electron microscopy [4, 5, 38]. The tBFI maps distinctly reveal the terminal central venules with a diameter of 29.8 *±* 1.5*μ*m (mean *±* SEM, n=14 vessels from 3 rats). The sinusoids converging onto these vessels had a diameter of 10.1 *±* 0.5 *μ*m (mean *±* SEM, n=13 vessels from 2 rats) and can be clearly distinguished for up to 0.1–0.5 mm away from the large venules. Near the terminal central venules, the sinusoids run parallel to each other, while near the terminal portal venules they are tortuous (Fig. 1e,g; Fig. S2), revealling a typical morphology of pericentral and periportal sinusoids [8, 39]. The terminal portal venules visualize blurry in the tBFI maps because these vessels terminate beneath the liver capsule in rats [14, 20] and, hence, are out of focus. Additional blurring may result from static scattering caused by tissue overlying the terminal portal venules.

### 2.2 Angiotensin II decreases blood flow in hepatic sinusoids and venules

Angiotensin II is a vasoconstrictor known to reduce hepatic blood flow [12, 13]. Consistently, injections of Angiotensin II upstream of the portal vein reduced hepatic tBFI in sinusoids and terminal central venules (Fig. 2). As a control, we injected saline, which increased tBFI only for *<*40 seconds (Fig. 2a–c). The magnitude of the tBFI increase was larger at higher injection rates (Fig. 2c), suggesting that the increase in hepatic blood flow was caused by the injection volume. Angiotensin II injections triggered a biphasic response (Fig. 2d). First, tBFI increased similar to the response to the saline injection, hence, explained by the injection volume. Second, tBFI decreased, indicating a reduction of the hepatic blood flow consistent with the vasoconstrictive effect of Angiotensin II (Fig. 2d, top) [13]. The decrease in tBFI was only induced by larger doses of Angiotensin II (5 and 20 ng), but not smaller doses (1 and 3 ng). This response was not due to hypotension, as blood pressure increased as expected following Angiotensin II administration (Fig. 2d, bottom). These findings demonstrate the sensitivity of LSCI in detecting changes for hepatic blood flow dynamics induced by vasoconstrictive agents.

**Figure 2:**
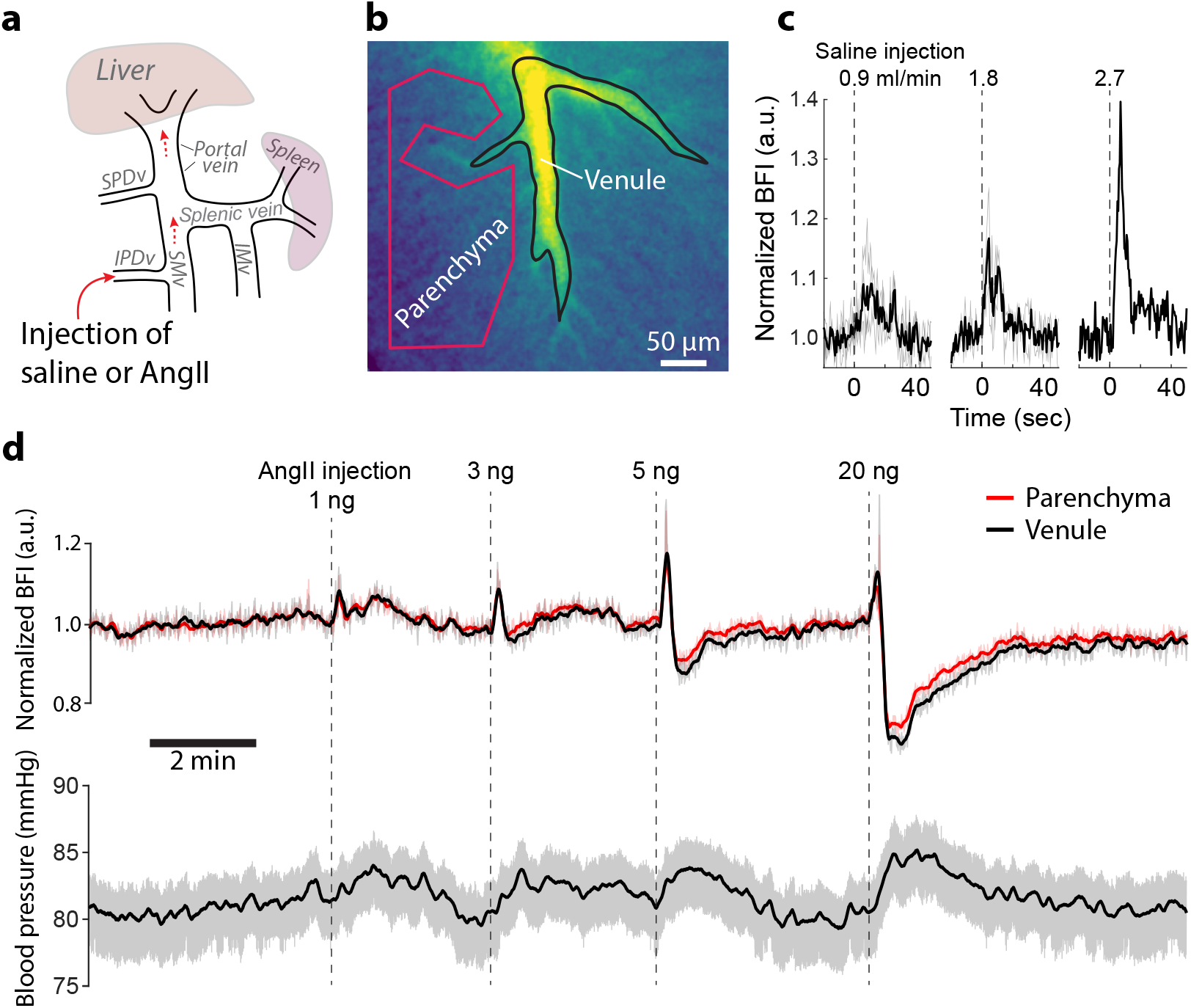
Response of hepatic superficial vessels to the local injection of saline and Angiotensin II. **a**) Major veins supplying blood to the liver. Dashed arrows indicate the direction of the blood flow. Saline or Angiotensin II were injected into the inferior pancreaticoduodenal vein (IPD v.). **b**) A tBFI map and regions of interest (ROIs) drawn over a terminal portal venule (black) and parenchyma which contains hepatic sinusoids (red). **c**) Responses of the normalized tBFI response in terminal central venule to saline (150 *μ*l) infused at different rates. The infusion started at time zero. Grey lines are individual recordings while black lines are averages of the grey lines (only for 0.9 and 1.8 ml/min). **d**) *Top*: response of the normalized tBFI from the venular (black) and the parenchyma (red) ROIs to Angiotensin II infusions at different doses (volume 150 *μ*l, injection rate 0.9 ml/min). The tBFI was normalized to the average tBFI before the first injection. Thin lines are the raw tBFI averaged within the ROIs, and thick lines are the smoothed thin lines. *Bottom*: arterial blood pressure during the recording. The black line is the smoothed raw blood pressure. Abbreviations: SPD v. — superior pancreaticoduodenal vein; IMv and SMv — inferior and superior mesenteric veins, respectively.

### 2.3 Oscillations of hepatic blood flow

Oscillations of blood flow play an important role in the brain, kidney and other organs [40–43]. However, their presence in the liver microcirculation remains ellusive. We observed that the spatial blood flow index (sBFI) oscillates in sinusoids, terminal portal venules, and terminal central venules under isoflurane and sevoflurane anesthesia (Fig. 3, Fig. S3). These oscillations occur consistently between sinusoids and venules situated up to 1 mm apart (Fig. 3a,b), suggesting that they originate from upstream blood vessels, including the portal vein and the hepatic artery. Temporally, the oscillations are intermittent (Fig. 3c), with frequencies falling within the 0.05–0.4 Hz range and varying between individual rats (Fig. S3a,b). These frequencies are consistent with arterial vasomotion (0.05–0.2 Hz) [43] and rhythmic contractions of the portal vein (0.07–0.2 Hz) [16]. Although isoflurane and sevoflurane may differently affect hepatic vascularure [44–50], the sBFI’s power within the vasomotion frequency interval was not significantly different between the two anesthetics (p ¡ 0.05, Mann-Whitney U test; Fig. S3c).

**Figure 3:**
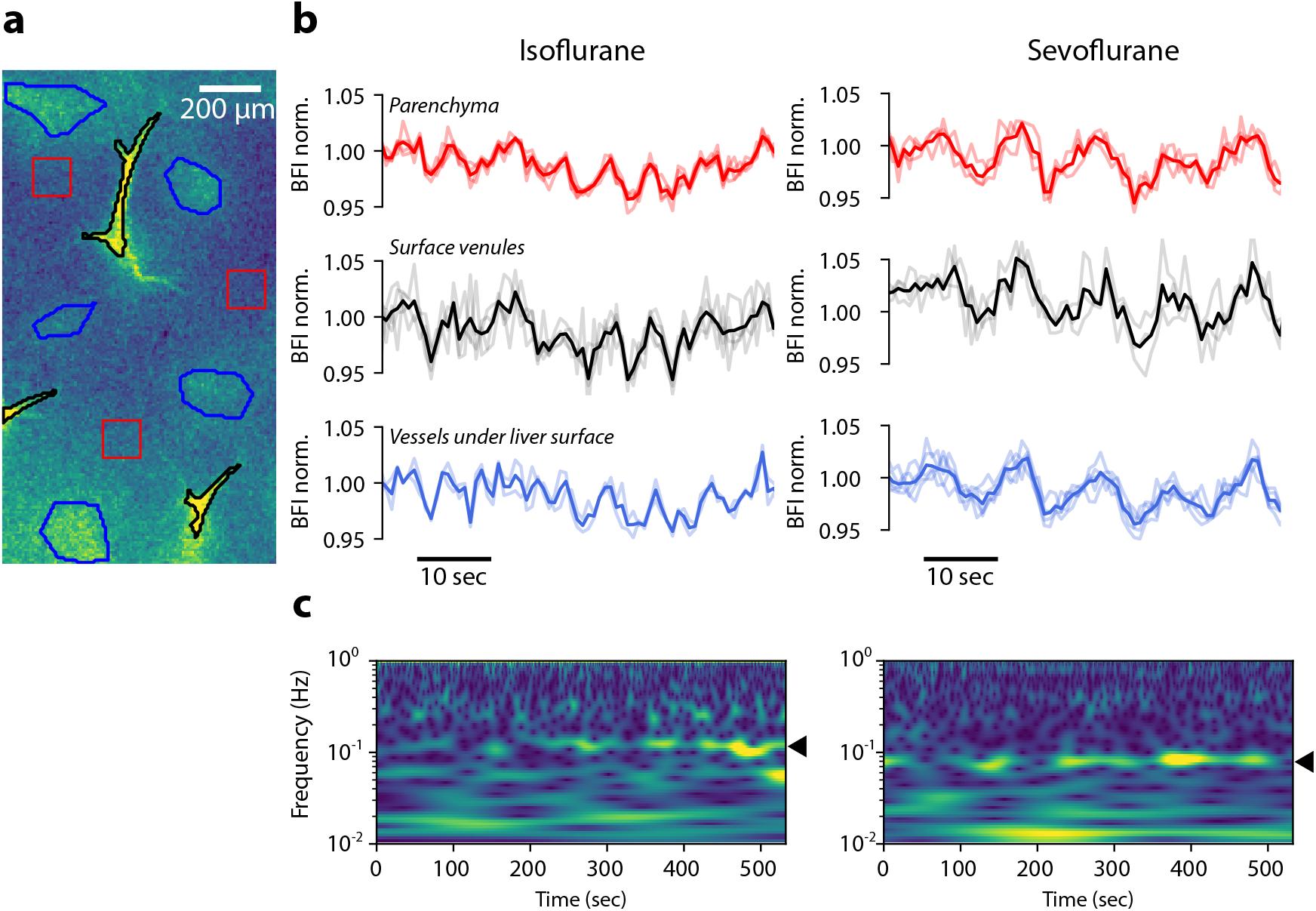
Dynamics of the hepatic blood flow observed under isoflurane and sevoflurane anesthesia. **a**) sBFI map with several regions of interest in parenchyma (red), venules of group 1 (black) and 2 (blue). **b**) Normalized sBFI in a rat anesthetized with isoflurane (*left*) and sevoflurane (*right*). Each transparent line is the Sbfi averaged in a single ROI and smoothed by averaging in non-overlapping blocks of 43 points (0.86 sec). The solid lines are averages over different ROIs. **c**) Wavelet spectrograms of the sBFI averaged over parenchyma and blood vessels in the rats anesthetized with isoflurane (*left*) and sevoflurane (*right*). The spectrograms show the distribution of periodic changes in the time-frequency domain. Arrowheads point to recurrent oscillations. The range of the magnitudes of the wavelet transform is the same in both spectrograms. See also Fig. S3a,b showing power spectral densities from all rats.

## 3 Discussion

In this study, we demonstrated the utility of LSCI for investigation of the liver hemodynamics at the microvascular level *in vivo*. Our findings show that LSCI spatially resolves individual hepatic blood vessels — sinusoids, terminal central venules, and terminal portal venules — and it is sensitive to reductions in liver blood flow induced by local adminisstration of Angiotensin II and to spontaneous oscillations of the hepatic blood flow.

Previous LSCI studies did not achieve vessel-specific blood flow measurements in the liver possibly due to large image pixel sizes (about 100 *μ*m/pixel [34, 51]), or motion blur [33–35, 51]. In contrast, our LSCI images have a finer pixel size (1.19 *μ*m/pixel), which, combined with liver stabilization and image registration, enabled resolution of individual hepatic microvessels. Our estimates of the diameter of terminal central venules (29.8 *±* 1.5*μ*m) and pericentral sinusoids (10.1 *±* 0.5 *μ*m) are similar to previous measurements [19, 36, 52].

We observed that blood flow oscillates in hepatic terminal portal venules, sinusoids, and terminal central venules at frequencies consistent with vasomotion. A possible explanation is vasomotion of the hepatic artery (0.05–0.2 Hz) [43] and rhythmic contractions of the portal vein (0.07–0.2 Hz) [16]. Supporting this hypothesis, we found that the sBFI oscillations were similar in different liver microvessels. While isoflurane and sevoflurane can inhibit the arterial vasomotion by vasodilation [42, 53–56], the effect of these anesthetics on the rhythmic contraction of the portal vein remains unclear [49, 57–59]. It is possible that the portal vein contractions cause the observed flowmotion in hepatic microvessels, but more research is needed to test this hypothesis.

We propose that LSCI is a valuable tool to study the mechanisms underlying sinusoidal blood flow regulation, which is critical for liver function. Dysregulation of sinusoids is increasingly linked to portal hypertension [60], metabolic dysfunction-associated steatotic liver disease [61–63], liver metastases [64], and other pathologies [65]. Furthermore, recent advances in understanding of molecular zonation of hepatocytes [66–68] raise questions about how sinusoids balance blood flow between different hepatocyte populations. Sinusoids can regulate flow by changing their diameter [19, 52], possibly via contractions of the hepatic stellate cells and by constriction of sinusoidal sphincters [20, 69–71], but the relationship to liver pathology remains obscure. LSCI offers an oportunity to study the regulation of sinusoidal blood flow in liver diseases over large fields of view *in vivo*.

Our study has several limitations. First, while LSCI provides horizontal resolution of blood vessles, it integrates scattered light from a layer of tissue likely containing several vessels. In the brain, the penetration depth of LSCI was estimated to be 50–150 *μ*m [29, 72, 73], but in the liver such estimate does not exist yet. Consequently, the BFI measured in the terminal central and portal venules may be affected by overlaying/underlying sinusoids. In parenchyma, the BFI corresponds to the blood flow in a population of sinusoids rather than individual ones. A second limitation is that our liver stabilization was not perfect, and some respiration-induced blurring of speckle contrast compromized the data quality. Additional stabilization of the liver could be acheived by submerging the liver under agarose gel, as demostrated previously [74]. Lastly, our study was done in rats anesthetized with isoflurane or sevoflurane, which can alter the portal venous flow, although maintaining overall liver perfusion [44, 47–50, 59].

## 4 Methods

### 4.1 Animal preparation

All experimental procedures were approved by the Danish National Committee on Health Research Ethics in accordance with the EU Directive 2010/63/EU for animal experiments. This study used healthy Sprague-Dawley rats (N=11 males, Janvier Labs, France) housed with 12/12 hour light/dark cycle and ad-libitum access to chow and water.

We performed preparatory surgery as described previously in Ref. [30]. Briefly, the rats were anestehtized with isoflurane (5 % induction and surgery, 1.5 % imaging, n=6 rats) or sevoflurane (8 % induction, 5 % surgery, 2.5 % imaging, n=5 rats) in 35 % oxygen and 65 % nitrogen. The left common carotid artery and femoral vein were cannulated, respectively, for arterial blood pressure monitoring and infusion of cisatracurium (Nimbex®, 0.25 mg/ml in saline, 40 *μ*l/min) for muscle relaxation during mechanical ventilation. The trachea was intubated, and anesthesia delivery was changed from a mask to the trachea tube with mechanical ventilation. The body temperature was maintained by a servo-controlled heating table set up at 37 °C. The arterial blood pressure during LSCI recordings was (mean *±* SEM) 107 *±* 4 mmHg (n=5, sevoflurane), and 79 *±* 3 mmHg (n=5, isoflurane).

For imaging, the left lateral lobe of the liver was exposed and stabilized between two holders: one supported the liver from below and reduced the vertical movement; the other separated the liver from the diaphragm and reduced the lateral movement (Fig. 1a). A motion marker (bent 40 *μ*m-diameter wire) was placed on the liver surface to allow correction of the residual movement by image registration.

The effect of Angiotensin II on the hepatic blood flow was assessed in one rat anesthetized with isoflurane. 150 *μ*l of saline or Angiotensin II (Sigma-Aldrich, #05-23-0101; 1, 3, 5, and 20 ng in saline) was injected via a cannula placed in the inferior pancreaticoduodenal vein (IPD v.) (Fig. 2a). The injections were performed with the rates of 0.9, 1.8, and 2.7 ml/min (saline) and 0.9 ml/min (Angiotensin II) using a pump (AL-1000, World Precision Instruments).

### 4.2 Laser speckle imaging

Our imaging setup was previously described in detail in Ref. [74]. In brief, the liver tissue was illuminated with a holographic volume grating stabilized laser diode (785 nm, Thorlabs LP7850-SAV50) controlled by CLD101x (Thorlabs). The scattered light passed through a linear polarizing filter and was captured by a CMOS camera (Basler acA2048-90umNIR, 5.5 *μ*m pixel size, 8 bit mode). The camera was mounted on a VZM 450 lens with adjustable zoom (Edmund optics). The linear polarizing filter was adjusted to minimize the mean intensity of the speckle image to prevent capturing the reflected light.

The laser speckle images were recorded from 1.9*×*1.9-mm (1600*×*1600 pixels, 1.19 *μ*m/pixel) field of view at 50 Hz frame rate with 5 ms exposure time [29, 74]. Before the acquisition, the focus was adjusted such that the blood vessels on the liver surface, seen on an online-calculated spatial laser speckle contrast image, were as sharp as possible. The adjustable zoom of the lens allowed acquiring images from a larger field of view to capture overview of the liver microvasculature.

### 4.3 Calculation of laser speckle contrast and blood flow index

The speckle images were registered by calculating the translation (imregister command, MATLAB) of the movement marker in the Gaussian-blurred (sigma = 7 pixels) frames, and translating the raw speckle images according to the translations. The laser speckle contrast (LSC) was calculated from the registered images as the standard deviation divided by the mean of pixel values in temporal or spatial domain, depending on the application [75]. To preserve the spatial detail of the liver vasculature, we computed the temporal LSC (tLSC) in non-overlapping blocks of 25 frames (0.5 sec). To analyze the blood flow dynamics, we computed the spatial LSC (sLSC) in non-overlapping blocks of 7-by-7 pixels (8.3-by-8.3 *μ*m). Unlike tLSC, sLSC preserves the original frame rate of the data but reduces the spatial detail, obscurring individual sinusoids, yet distinguishing individual surface venules. As our primary interest is the hepatic blood flow, tLSC and sLSC were converted to the blood flow index (tBFI and sBFI, respectively), as the inverse squared contrast value [28, 74]. tBFI was then used to obtain tBFI maps of liver microvasculature, by temporal averaging, and to assess blood flow responses to injections of saline and Angiotensin II. sBFI was used to characterize spontaneous dynamics of the blood flow.

To estimate the changes of tBFI (sBFI) within sinusoids, terminal central venules, and terminal portal venules, we averaged the pixel values within regions of interest (ROIs) drawn manually over respective vessel types. The terminal central venules were identified as the large-diameter vessels in focus on tBFI (sBFI) images; the blood in this vessels flows from small-to large-diameter segments as we observed in a binocular microscope prior to LSCI. The terminal portal venules were identified as regions of the time-averaged tBFI (sBFI) images with high intensity pixels, without sharp edges. These vessels are found underneath the liver surface in rats [20] and, hence, are out-of-focus in our images. It was impossible to adjust the focus to obtain sharp images of the terminal portal venules likely due to the presence of static scattering from the tissue above the terminal portal venules. The sinusoids are located throughout the liver between the terminal central and portal venules [4, 8].

### 4.4 Estimation of vessel diameter

The diameter of the terminal central venules and sinusoids was estimated from the tBFI maps. For that, we obtained a one-dimensional profile across a vessel by (a) finding a straight segment of this vessel, (b) drawing its centerline, (c) defining a rectangle area containing the vessel segment, (d) averaging the pixel values within the rectangle in the direction of the centerline. In the profile, the vessel visualizes as a single peak. We estimated the vessel diameter by calculating the full width at half maximum (FWHM) of the peak. The diameter was estimated in 14 terminal central venules (n=3 rats) and in 11 sinusoids converging to terminal central venules (n=2 rats). For each vessel, we averaged up to 5 FWHM values measured in different segments.

### 4.5 Data analysis in frequency domain

The one-sided power spectral density (PSD) was estimated from the sBFI (averaged within parenchyma ROIs) by the Welch’s method (`welch` command, Scipy version 1.11.4, Python) with the window = 2^12^ points (81.92 sec), overlap = 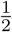 window, and the other parameters having their default values. The PSD was then averaged over ROIs (3–5 ROIs per rat). The stability of sBFI oscillations in time was assessed by the continuous wavelet transform (`cwt` command, MATLAB) of the sBFI computed with a generalized Morse wavelet with standard parameters (symmetry parameter, *γ* = 3 and time-bandwidth product = 60) [76].

## Funding

This study was supported by the Novo Nordisk Foundation (NNF22OC0079295).

## Disclosures

The authors declare no conflicts of interest.

## Data availability statement

Data underlying the results presented in this paper are not publicly available at this time but may be obtained from the authors upon reasonable request.

## Supplementary figures

**Figure S1:**
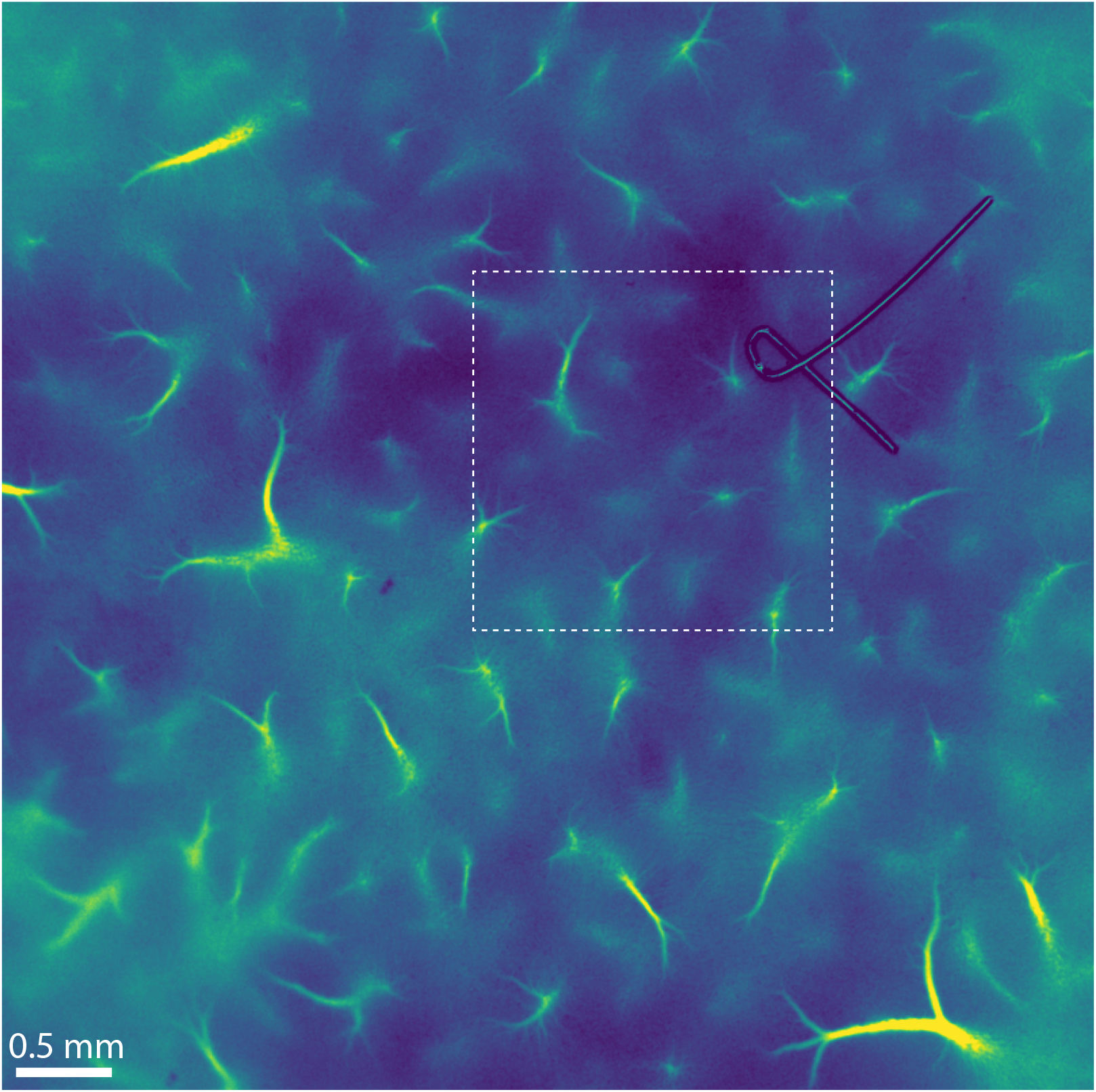
Overview of the hepatic microvasculature seen in a blood flow index map. Laser speckle data were recorded with 2*×* zoom set on the lens (Methods). The region inside the white rectangle was then imaged at 4.5*×* zoom (Fig. S2)

**Figure S2:**
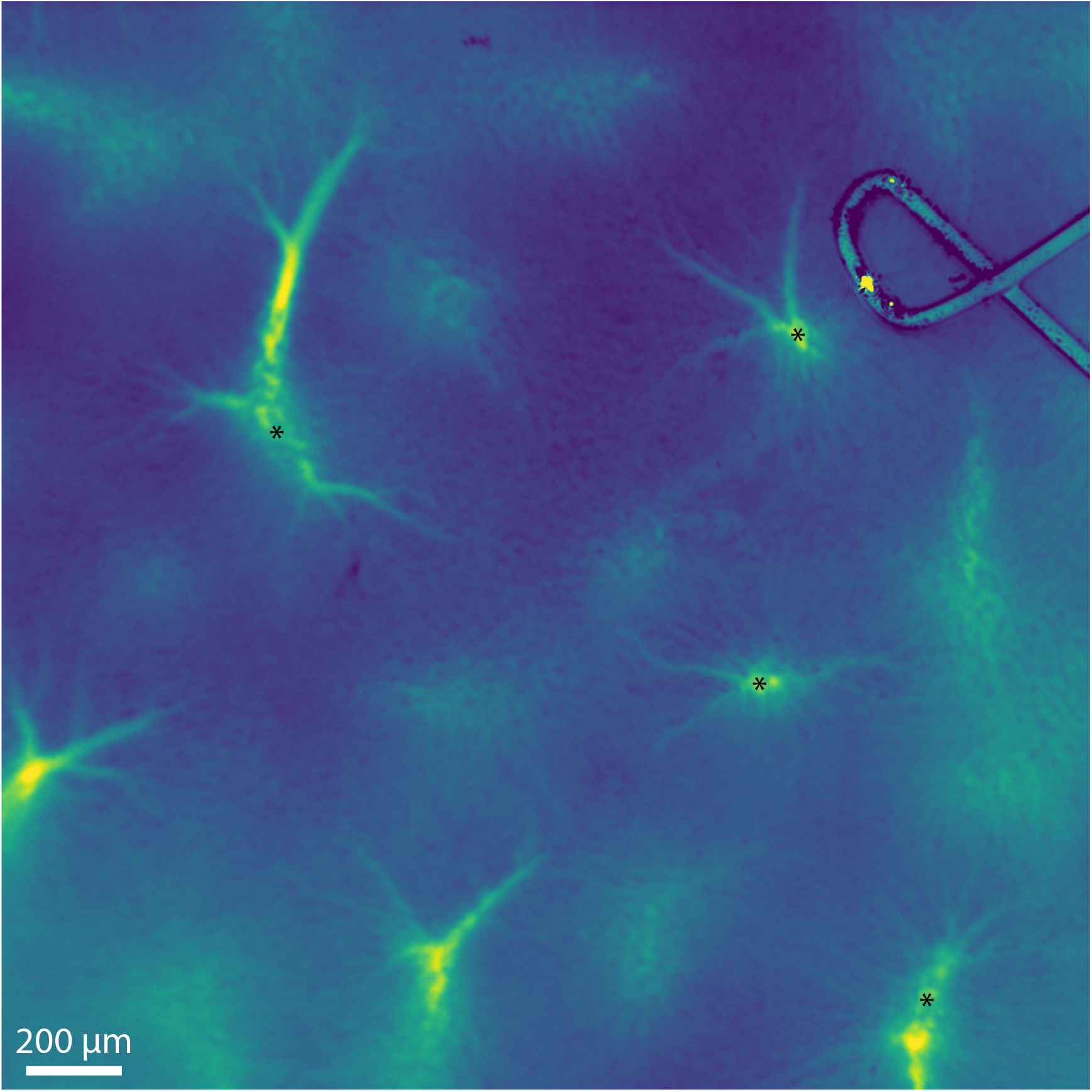
Blood flow index map of hepatic microvasculature obtained from a region in Fig. S1). Sinusoids can be seen clearly converging parallel to each other around some of the terminal central venules (asterisks).

**Figure S3:**
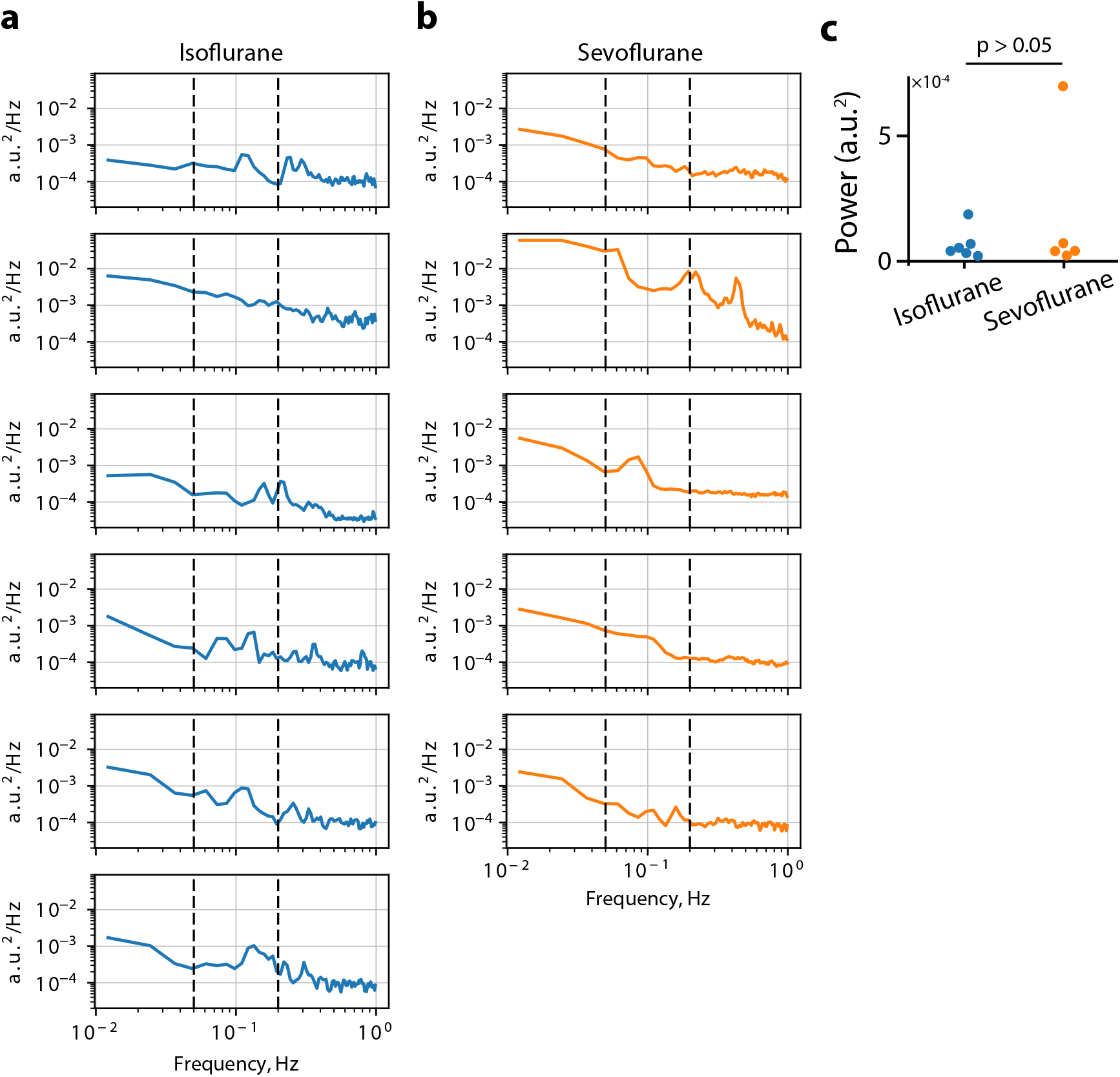
**a,b**) Power spectral density (PSD) of the blood flow index in parenchyma of different rats anesthetized with isoflurane (a) and sevoflurane (b). Each panel represents a single rat. The PSD was averaged over the ROIs (3-5 ROIs per rat). c) The signal’s power within the frequency range of 0.05-0.2 Hz in isoflurane-and sevoflurane-anesthetized rats (Mann-Whitney U test).

## Notes

### Competing Interest Statement

The authors have declared no competing interest.

### Summary of Updates

All text was improved for clarity. Discussion paragraphs were rearranged.

## References

1. Ozougwu JC. Physiology of the liver. International Journal of Research in Pharmacy and Biosciences 2017;4:13–24.

2. Eipel C, Abshagen K, and Vollmar B. Regulation of hepatic blood flow: the hepatic arterial buffer response revisited. World J. Gastroenterol. 2010;16:6046–57.

3. Greenway CV and Stark RD. Hepatic vascular bed. Physiol. Rev. 1971;51:23– 65.

4. Ohtani O and Murakami T. Peribiliary portal system in the rat liver as studied by the injection replica scanning electron microscope method. In: Scanning electron microscopy. USA: SEM Inc., AMF O’Hare, 1978:241.

5. Ohtani O, Murakami T, and Jones AL. Microcirculation of the liver, with special reference to the peribiliary portal system. In: Basic and Clinical Hepatology. Dordrecht: Springer Netherlands, 1982:85–96. doi: 10.1007/978-94-009-8216-1/_6.

6. Lametschwandtner A, Spornitz U, and Minnich B. Microvascular anatomy of the non-lobulated liver of adult Xenopus laevis: A scanning electron microscopic study of vascular casts. Anat. Rec. (Hoboken) 2022;305:243–53.

7. Trefts E, Gannon M, and Wasserman DH. The liver. Curr. Biol. 2017;27:R1147– R1151.

8. McCuskey RS. The hepatic microvascular system in health and its response to toxicants. Anat. Rec. (Hoboken) 2008;291:661–71.

9. Lautt WW. Mechanism and role of intrinsic regulation of hepatic arterial blood flow: hepatic arterial buffer response. Am. J. Physiol. 1985;249:G549–56.

10. Richter S, Vollmar B, Mücke I, Post S, and Menger MD. Hepatic arteriolo-portal venular shunting guarantees maintenance of nutritional microvascular supply in hepatic arterial buffer response of rat livers. J. Physiol. 2001;531:193– 201.

11. Mücke I, Richter S, Menger MD, and Vollmar B. Significance of hepatic arterial responsiveness for adequate tissue oxygenation upon portal vein occlusion in cirrhotic livers. Int. J. Colorectal Dis. 2000;15:335–41.

12. Pereira AJ, Jeger V, Fahrner R, et al. Interference of angiotensin II and enalapril with hepatic blood flow regulation. Am. J. Physiol. Gastrointest. Liver Physiol. 2014;307:G655–63.

13. Edgarian H and Altura BM. Ethanol and contraction of venous smooth muscle. Anesthesiology 1976;44:311–7.

14. Adori C, Daraio T, Kuiper R, et al. Disorganization and degeneration of liver sympathetic innervations in nonalcoholic fatty liver disease revealed by 3D imaging. Sci. Adv. 2021;7:eabg5733.

15. Robinson KA, Middleton WD, AL-Sukaiti R, Teefey SA, and Dahiya N. Doppler sonography of portal hypertension. Ultrasound Q. 2009;25:3–13.

16. Spencer NJ and Greenwood IA. Characterization of properties underlying rhythmicity in mouse portal vein. Auton. Neurosci. 2003;104:73–82.

17. Sironi L, Bouzin M, Inverso D, et al. In vivo flow mapping in complex vessel networks by single image correlation. Sci. Rep. 2014;4:7341.

18. Timm F and Vollmar B. Heterogeneity of the intrahepatic portal venous blood flow: impact on hepatocyte transplantation. Microvasc. Res. 2013;86:34–41.

19. Oda, M., Azuma, T., Watanabe, N., Nishizaki, Y., Nishida, J., Ishii, K., Suzuki, H., Kaneko, H., Komatsu, H., Tsukada, N., Tsuchiya M. Regulatory mechanisms of the hepatic microcirculation-involvement of the contraction and dilatation of sinusoids and sinusoidal endothelial fenestrae. In: Progress in Applied Microcirculation; vol. 17. Ed. by K. Messmer H and F. Hammersen M. Basel, Switzerland: S Karger AG, 1990.

20. Oda M, Yokomori H, and Han JY. Regulatory mechanisms of hepatic microcirculation. Clin. Hemorheol. Microcirc. 2003;29:167–82.

21. Li DY, Xia Q, Yu TT, Zhu JT, and Zhu D. Transmissive-detected laser speckle contrast imaging for blood flow monitoring in thick tissue: from Monte Carlo simulation to experimental demonstration. Light Sci. Appl. 2021;10:241.

22. Phan T, Crouzet C, Kennedy GT, Durkin AJ, and Choi B. Quantitative hemo-dynamic imaging: a method to correct the effects of optical properties on laser speckle imaging. Neurophotonics 2023;10:045001.

23. Heeman W, Steenbergen W, Dam G van, and Boerma EC. Clinical applications of laser speckle contrast imaging: a review. J. Biomed. Opt. 2019;24:1–11.

24. Briers D, Duncan DD, Hirst E, et al. Laser speckle contrast imaging: theoretical and practical limitations. J. Biomed. Opt. 2013;18:066018.

25. Boas DA and Dunn AK. Laser speckle contrast imaging in biomedical optics. J. Biomed. Opt. 2010;15:011109.

26. Dunn AK, Huang Z, Boas DA, and Moskowitz MA. Intrinsic brain activity triggers trigeminal meningeal afferents in a migraine model. Nat. Med. 2002.

27. Mikkelsen SH, Skøtt MV, Gutierrez E, and Postnov DD. Laser speckle imaging of the hippocampus. Biomed. Opt. Express 2024;15:1268.

28. Holstein-Rathlou NH, Sosnovtseva OV, Pavlov AN, Cupples WA, Sorensen CM, and Marsh DJ. Nephron blood flow dynamics measured by laser speckle contrast imaging. Am. J. Physiol. Renal Physiol. 2011;300:F319–29.

29. Postnov D, Marsh DJ, Cupples WA, Holstein-Rathlou NH, and Sosnovtseva O. Synchronization in renal microcirculation unveiled with high-resolution blood flow imaging. Elife 2022;11.

30. Lee B, Postnov DD, Sørensen CM, and Sosnovtseva O. The assessment of cortical hemodynamic responses induced by tubuloglomerular feedback using in vivo imaging. Physiol Rep 2023;11:e15648.

31. Heeman W, Maassen H, Calon J, et al. Real-time visualization of renal microperfusion using laser speckle contrast imaging. J. Biomed. Opt. 2021;26:056004.

32. Neganova AY, Postnov DD, Sosnovtseva O, and Jacobsen JCB. Rat retinal vasomotion assessed by laser speckle imaging. PLoS One 2017;12:e0173805.

33. Sturesson C, Milstein DMJ, Post ICJH, Maas AM, and Gulik TM van. Laser speckle contrast imaging for assessment of liver microcirculation. Microvasc. Res. 2013;87:34–40.

34. Li CH, Wang HD, Hu JJ, et al. The monitoring of microvascular liver blood flow changes during ischemia and reperfusion using laser speckle contrast imaging. Microvasc. Res. 2014;94:28–35.

35. Li CH, Ge XL, Pan K, Wang PF, Su YN, and Zhang AQ. Laser speckle contrast imaging and Oxygen to See for assessing microcirculatory liver blood flow changes following different volumes of hepatectomy. Microvasc. Res. 2017;110:14– 23.

36. Kan Z and Madoff DC. Liver anatomy: microcirculation of the liver. Semin. Intervent. Radiol. 2008;25:77–85.

37. Bloch EH. The in vivo microscopic vascular anatomy and physiology of the liver as determined with the quartz rod method of transillumination. Angiology 1955;6:340–9.

38. Grisham JW, Nopanitaya W, Compagno J, and Nägel AE. Scanning electron microscopy of normal rat liver: the surface structure of its cells and tissue components. Am. J. Anat. 1975;144:295–321.

39. Hase T and Brim J. Observation on the microcirculatory architecture of the rat liver. Anat. Rec. 1966;156:157–73.

40. Lee B, Postnov DD, Sørensen CM, and Sosnovtseva O. In vivo mapping of hemodynamic responses mediated by tubuloglomerular feedback in hypertensive kidneys. Sci. Rep. 2023;13:21954.

41. Broggini T, Duckworth J, Ji X, et al. Long-wavelength traveling waves of vasomotion modulate the perfusion of cortex. Neuron 2024;112:2349–2367.e8.

42. Veluw SJ van, Hou SS, Calvo-Rodriguez M, et al. Vasomotion as a driving force for paravascular clearance in the awake mouse brain. Neuron 2020;105:549–561.e5.

43. Aalkjær C, Boedtkjer D, and Matchkov V. Vasomotion - what is currently thought? Acta Physiol. 2011;202:253–69.

44. Limmen J van, Wyffels P, Berrevoet F, et al. Effects of propofol and sevoflurane on hepatic blood flow: a randomized controlled trial. BMC Anesthesiol. 2020;20:241.

45. Gelman S. General anesthesia and hepatic circulation. Can. J. Physiol. Pharmacol. 1987;65:1762–79.

46. Frink E. The hepatic effects of sevoflurane. Anesth. Analg. 1995;81:S46–50.

47. Jacob L, Boudaoud S, Payen D, et al. Isoflurane, and not halothane, increases mesenteric blood flow supplying esophageal ileocoloplasty. Anesthesiology 1991;74:699–704.

48. Grundmann U, Zissis A, Bauer C, and Bauer M. In vivo effects of halothane, enflurane, and isoflurane on hepatic sinusoidal microcirculation. Acta Anaesthesiol. Scand. 1997;41:760–5.

49. Wilde DW. Isoflurane reduces K+ current in single smooth muscle cells of guinea pig portal vein. Anesth. Analg. 1996;83:1307–1313.

50. Gatecel C, Losser MR, and Payen D. The postoperative effects of halothane versus isoflurane on hepatic artery and portal vein blood flow in humans. Anesth. Analg. 2003;96:740–5.

51. Eriksson S, Nilsson J, Lindell G, and Sturesson C. Laser speckle contrast imaging for intraoperative assessment of liver microcirculation: a clinical pilot study. Med. Devices 2014;7:257–61.

52. Zhang JX, Pegoli Jr W, and Clemens MG. Endothelin-1 induces direct constriction of hepatic sinusoids. Am. J. Physiol. 1994;266:G624–32.

53. Tanaka K, Kawano T, Nakamura A, et al. Isoflurane activates sarcolemmal adenosine triphosphate-sensitive potassium channels in vascular smooth muscle cells: a role for protein kinase A. Anesthesiology 2007;106:984–91.

54. Iida H, Ohata H, Iida M, Watanabe Y, and Dohi S. Isoflurane and sevoflurane induce vasodilation of cerebral vessels via ATP-sensitive K+ channel activation. Anesthesiology 1998;89:954–60.

55. Auer LM and Gallhofer B. Rhythmic activity of cat pial vessels in vivo. Eur. Neurol. 1981;20:448–68.

56. Colantuoni A, Bertuglia S, and Intaglietta M. Effects of anesthesia on the spontaneous activity of the microvasculature. Int. J. Microcirc. Clin. Exp. 1984;3:13–28.

57. Conzen P, Vollmar B, Habazettl H, Frink E, Peter K, and Messmer K. Systemic and regional hemodynamics of isoflurane and sevoflurane in rats. Anesth. Analg. 1992;74:79–88.

58. Frink Jr EJ, Morgan SE, Coetzee A, Conzen PF, and Brown Jr BR. The effects of sevoflurane, halothane, enflurane, and isoflurane on hepatic blood flow and oxygenation in chronically instrumented greyhound dogs. Anesthesiology 1992;76:85–90.

59. Bernard JM, Doursout MF, Wouters P, Hartley CJ, Merin RG, and Chelly JE. Effects of sevoflurane and isoflurane on hepatic circulation in the chronically instrumented dog. Anesthesiology 1992;77:541–5.

60. Gracia-Sancho J, Marrone G, and Fernández-Iglesias A. Hepatic microcirculation and mechanisms of portal hypertension. Nat. Rev. Gastroenterol. Hepatol. 2019;16:221–34.

61. Terkelsen MK, Bendixen SM, Hansen D, et al. Transcriptional dynamics of hepatic sinusoid-associated cells after liver injury. Hepatology 2020;72:2119– 33.

62. Graaff D van der, Kwanten WJ, and Francque SM. The potential role of vascular alterations and subsequent impaired liver blood flow and hepatic hypoxia in the pathophysiology of non-alcoholic steatohepatitis. Med. Hypotheses 2019;122:188–97.

63. Bendixen SM, Jakobsgaard PR, Hansen D, et al. Single cell-resolved study of advanced murine MASH reveals a homeostatic pericyte signaling module. J. Hepatol. 2024;80:467–81.

64. Tsilimigras DI, Brodt P, Clavien PA, et al. Liver metastases. Nat. Rev. Dis. Primers 2021;7:27.

65. Gracia-Sancho J, Caparrós E, Fernández-Iglesias A, and Francés R. Role of liver sinusoidal endothelial cells in liver diseases. Nat. Rev. Gastroenterol. Hepatol. 2021;18:411–31.

66. Jungermann K and Katz N. Functional specialization of different hepatocyte populations. Physiol. Rev. 1989;69:708–64.

67. Paris J and Henderson NC. Liver zonation, revisited. Hepatology 2022;76:1219– 30.

68. Halpern KB, Shenhav R, Matcovitch-Natan O, et al. Single-cell spatial reconstruction reveals global division of labour in the mammalian liver. Nature 2017;542:352–6.

69. McCuskey RS. A dynamic and static study of hepatic arterioles and hepatic sphincters. Am. J. Anat. 1966;119:455–77.

70. Burkel WE. The fine structure of the terminal branches of the hepatic arterial system of the rat. Anat. Rec. 1970;167:329–49.

71. Knisely MH, Harding F, and Debacker H. Hepatic sphincters; brief summary of present-day knowledge. Science 1957;125:1023–6.

72. Davis MA, Kazmi SMS, and Dunn AK. Imaging depth and multiple scattering in laser speckle contrast imaging. J. Biomed. Opt. 2014;19:086001.

73. Zheng S, Xiao S, Kretsge L, Cruz-Martín A, and Mertz J. Depth resolution in multifocus laser speckle contrast imaging. Opt. Lett. 2021;46:5059–62.

74. Lee B, Sosnovtseva O, Sørensen CM, and Postnov DD. Multi-scale laser speckle contrast imaging of microcirculatory vasoreactivity. Biomed. Opt. Express 2022;13:2312– 22.

75. González Olmos A, Humlesen Z, Matchkov V, and Postnov DD. Lossless temporal contrast analysis of laser speckle images from periodic signals. Biomed. Opt. Express 2023;14:1355–63.

76. The MathWorks Inc. Wavelet Toolbox version: 9.4 (R2024b). Natick, Massachusetts, United States, 2024. url: https://se.mathworks.com/products/wavelet.html.

